# Covering Hierarchical Dirichlet Mixture Models on binary data to enhance genomic stratifications in Onco-Hematology

**DOI:** 10.1101/2023.06.26.546639

**Authors:** Daniele Dall’Olio, Eric Sträng, Amin T Turki, Jesse M Tettero, Martje Barbus, Renate Schulze-Rath, Javier Martinez Elicegui, Tommaso Matteuzzi, Alessandra Merlotti, Luciana Carota, Claudia Sala, Matteo G Della Porta, Enrico Giampieri, Jesús María Hernández-Rivas, Lars Bullinger, Gastone Castellani, HARMONY Healthcare Alliance Consortium

## Abstract

Onco-hematological studies are increasingly adopting statistical mixture models to support the advancement of the genetically-driven classification systems for blood cancer. Targeting enhanced patients stratification based on the sole role of molecular biology attracted much interest and contributes to bring personalized medicine closer to reality. In particular, Dirichlet processes have become the preferred method to approach the fit of mixture models. Usually, the multinomial distribution is at the core of such models. However, despite their advanced statistical formalism, these processes are not to be considered black box techniques and a better understanding of their working mechanisms enables to improve their employment and explainability. Focused on genomic data in Acute Myeloid Leukemia, this work unfolds the driving factors and rationale of the Hierarchical Dirichlet Mixture Models of multinomials on binary data. In addition, we introduce a novel approach to perform accurate patients clustering via multinomials based on statistical considerations. The newly reported adoption of the Multivariate Fisher’s Non-Central Hypergeometric distributions reveals promising results and outperformed the multinomials in clustering both on simulated and real onco-hematological data.

**Author summary:** Explainable models are particularly attractive nowadays since they have the advantage to convince clinicians and patients. In this work we show that a deeper understanding of the Hierarchical Dirichlet Mixture Model, a non-black box method, can lead to better data modelling. In onco-hematology Hierarchical Dirichlet Mixture Models typically help to cluster molecular alterations rather than patients. Here, an intuitive statistical approach is presented to tackle patient classification based on the Hierarchical Dirichlet Mixture Models outcome. Additionally, molecular alterations are usually modelled by Hierarchical Dirichlet Mixture Models as a mixture of multinomial distributions. This work highlights that the alternative Fisher’s Non-Central Hypergeometric distribution can provide even better results and can give a higher priority to rare molecular alterations for patient classification.

## Introduction

In medicine, onco-hematology is leading the way towards personalized medicine thanks to the efforts dedicated to characterize and to cluster the genomic nature of blood tumors. Clustering is a paramount task carried out in several fields of data-driven science [1]. The ability to organize data in meaningful groups opens the possibility to identify shared characteristics that bring observations together and discriminate others that push them apart. Better knowledge of such characteristics contributes to better profile observations and to draw observation-specific considerations, i.e., personalization. In onco-hematology the object of clustering is usually the genomic information representing the presence or absence of genomic alterations, i.e., typically gene mutations and karyotypic anomalies.

To perform clustering nonparametric Bayesian methods are becoming of paramount importance in several field of medicine, including cancer research [2]. Especially with categorical data Hierarchical Dirichlet Mixture Models (HDMMs), which are notorious nonparametric Bayesian methods, are quickly gauging interest [3–7]. Their usage already proved to boost the definition of new clinical classification systems and the discovery of unknown groups of co-occurrent genomic alterations. The HDMMs have also been used transversally on genomic data, including copy number variation plus transcriptomic integration [8], pan-cancer proteomic characterization [9], cancer subtyping with microRNA [10] and disease classes discrimination based on genomics, transcriptomics and epigenomics [11].

In this paper we address the usage of the HDMMs in onco-hematology, due to the recent pioneering contributions in this field. Although the ultimate goal is to cluster patients, the application of HDMMs usually follows three steps: (a) a complete binary matrix of patients and genomic features is collected and pre-processed, (b) one or several HDMMs are run to detect groups of genomic features, (c) these groups are used to support a clinically-tailored definition of clusters for patients. The autonomous ability of HDMMs to determine the number of groups of genomic features and their direct interpretability are the main appealing reasons of the recent increased utilization of HDMMs. Given this increasing attention and usage, here, we reviewed methodologically the theory behind HDMMs along with results focused on novel and intuitive developments both on how to improve the prioritization of genomic features in groups and on how to statistically perform and generalize patients clustering. In the following sections the framework of the HDDM-based approaches is reported.

### Mixture models

Mixture models enable to model observations as a combination of multiple distributions [12]. An observation *x*_*n*_ for a sample *X*_*n*_ is usually supposed to be drawn from a single probability density function *f*(*X*_*n*_ = *x*_*n*_). Mixture models, instead, assume an observation to be sampled from *G* components (or groups) with prior probabilities {*π*_*g*_}_*g*=1..*G*_ and with conditional densities *f*_*g*_(*X*_*n*_ = *x*_*n*_|*π*_*g*_). The density function for sample *X*_*n*_ is the marginal density:

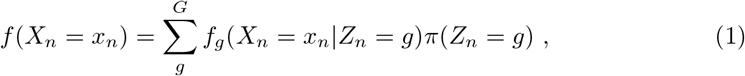

where the proxy variable *Z*_*n*_ indicates the component an observation is drawn from. Upon drawing from the mixture model, the proxy *Z*_*n*_ becomes a binary random variable that is zero for all component except being one for a single *g*-th. Therefore *Z*_*n*_ can be replaced by a random vector 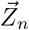 whose observations are single trials from a multinomial distribution with parameters equal to the prior component probabilities. That is,

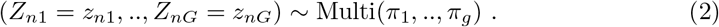

Thus, the sampling procedure from a mixture model first requires to draw a *g*-th component according to (2) and, secondly, it requires to sample from the corresponding density *f*_*g*_(·). In onco-hematology the widespread assumption is that patients’ genomic data derives from a mixture of multinomial components, so that the parameters of the density functions are probabilities.

### Dirichlet Mixture Model

Dirichlet Processes (DPs) [13] can be seen as the a priori processes of mixture models. The study of DPs usually involves other processes such as the Chinese Restaurant Process (CRP) [14] and the Blackwell-MacQueen (BM) models [15]. These processes fall outside the scope of this work but ultimately DP generalizes them both. Now, if a probability distribution *H* over the parameters of the components is defined, it is immediate to determine which parameters are more common, or which component is. Though, *H* is one among many potential distributions and its choice limit the exploration of the parameters space. What the DPs address is to provide a statistical framework to handle many possible a priori distributions. In other words, a DP is a probability distribution over the possible probability distributions for the parameters, which we generally refer to as *ϕ*_*k*_. Formally,

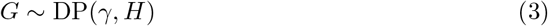

means that

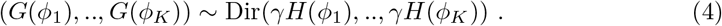

From a discrete perspective the above formulation implies that,

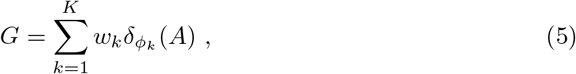

where the weights represent probabilities, i.e., *w*_*k*_ = *G*(*ϕ*_*k*_), and as such 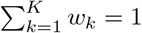 Since there is no restriction on *K* the number of components can be infinite (*K* =∞). Though, the number of components is never infinite, which implies that only a few components in the mixture have a non-zero probability of occurrence. This fact is usually described by processes with rich components that progressively get richer. All these concepts boil down to the complete formulation of the Dirichlet Mixture Model (DMM), which is described by:

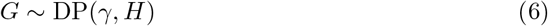

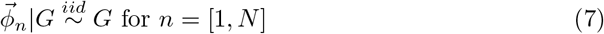

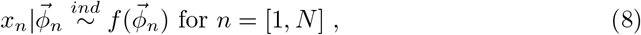

where iid stands for identically and independently distributed. In the onco-hematological scenario the HDMM intends to potentially cluster patients in infinite components, whose parameters are typical probabilities for the multinomials. During the Dirichlet Mixture Model (DMM) fit, the underlying “rich get richer” mechanism consents to automatically detect a probability distribution *G* of parameters. Since the majority of possible parameters have zero probability, i.e., *w*_*k*_ = 0, using a DP a priori gives the benefit of not choosing the number of expected components.

### Hierarchical Dirichlet Mixture Model

The DMM can be further extended by other DPs when observations are believed to be organized in groups [16]. Following the previous formulation, a realization from a Hierarchical Dirichlet Mixture Model (HDMM) follows:

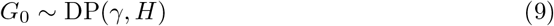

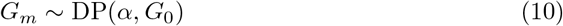

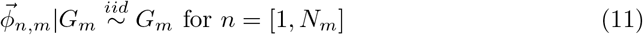

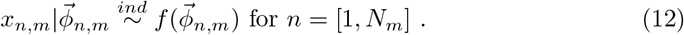

Typically, the modelling of onco-hematological genomic data is performed with a HDMM. In this case, the objects clustered by the HDMM are the single genomic alterations of each patient and not the whole patients with their co-existing sets of genetic alterations. With this perspective change each patient is considered as a separate group of genomic alterations. The main difference w.r.t. the DMM is that each patient is endowed with its own a priori probability distribution of parameters and an alteration is clustered also based on the patient it belongs to. Clearly, multiple levels can be added to the HDMM structure and the greater the number of levels (e.g. gender, age, etc.), the greater will be the sensitivity of the fit to retrieve group-related patterns.

## Materials and methods

### Extracting components

Once the statistical foundation of the HDMM is set and datasets with no-missing value are prepared, the HDMM needs to be fitted. Usually, Gibbs sampling [17] is employed and assigns, at each step of such Markov Chain Monte Carlo (MCMC) the observations, e.g. every genomic alteration in each patient, to a component one by one. Along the MCMC the number of components changes based on the observations they were assigned to in each previous step. Usually the MCMC is set to start from completely random components and the first iterations are excluded to first reach convergence. Upon convergence, every iteration along the MCMC estimates how observations are clustered and how many components exist. Hence, we obtained a sample from a HDMM as a matrix counting the number of times an observation is assigned to each component. Though, since many iterations compose a MCMC, we collected a matrix of this kind at every step. Additionally, we ran multiple MCMCs to address the dependency of the HDMM samples w.r.t. to the starting random sampling, which expands the collection of matrix samples. Eventually, we used all these matrix samples to determine the average number of times an observation belongs to every component. This approach is the standard one in onco-hematological genetic studies to determine groups (i.e., components) of genomic alterations and is based on the assumption that the data can be represented as a mixture of multinomial distributions, which shows how frequent each genomic alteration is in every component. To better adapt to the binary nature of the input data and to increase the importance of discriminative genomic alterations between components, we alternatively utilized the Fisher’s Non-Central Hypergeometric distribution (MFNCH) [18]. To our knowledge this is the first time the MFNCHs to better characterize clusters derived from the HDMM. Therefore, we eventually modelled the data as a mixture of MFNCH distributions. To estimate the MFNCH odds for each observation we adopted the Cornfield’s approximation [19] after estimating the observation mean (e.g. average occurences for a genomic alteration in a component).

### Data simulation

In total, 40 datasets were simulated that included one thousand data samples (*N*=1000) featuring fifty (*M*=50) binary events. Half of the datasets were drawn from five MFNCH distributions (*K* = 5), while the others from ten (*K* = 10). Plus, these MFNCH distributions were generated according to either a concentration parameter equal to one (*α*_*sim*_=1) or to 1*/M*. Datasets further differed based on the average number of observation per data samples (e.g. genomic alterations per patient) that spanned from one to ten. Each of the 40 datasets was modelled by two HDMMs: one driven by a concentration parameter *α*_*HDMM*_ of one or 1*/M*. Therefore 80 HDMMs were targeted and ten MCMCs were run for each of them to gather ten independent estimates, resulting in 800 running MCMCs.

### Classification approach

The HDMM, along with the chosen mixture models, determines only groups of observations, which are tipically used to build ad-hoc classifiers. Statistically, though, a whole data sample (e.g. patient) is assumed to be a product of the estimated mixture, i.e., it can originate potentially from every single component of the mixture. In order to define an intuitive and objective way to univocally assign a data sample to a component, we leveraged on the probability mass functions (p.m.f.) of either the multinomial or MFNCH distribution. In other words, once the *K* mixture components and their parameters, 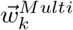 or 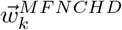, were estimated, we assign a data sample to the component with the highest p.m.f. value. With the multinomials scenario the assignment approach for each *i*-th data sample can be formulated as:

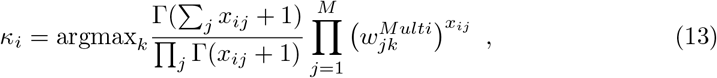

where *κ*_*i*_ is the component the i-th data sample is assigned to. The *x*_*ij*_ represents the value of the *j*-th observation of the *i*-th data sample. The same principle applies for MFNCH distribution, where the assignment formula becomes:

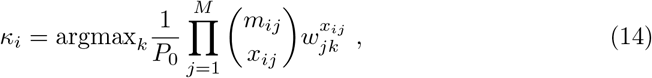

where *P*_0_ is the partition function and *m*_*ij*_ is the maximum number of times the *j*-th observation can potentially occur for the *i*-th data sample. Since all observations are binary, i.e., zero or one, all *m*_*ik*_ are set to one, and formula 14 simplifies to:

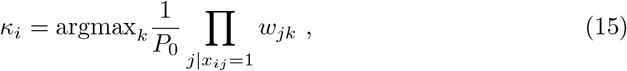

with

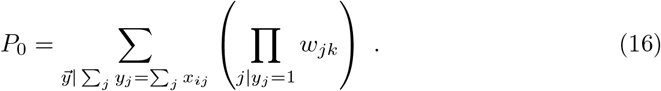

When multiple *k*s are associated to the maximum value, the data sample is considered ambiguous.

## Results

### HDMM convergence

The convergence of the 80 HDMMs was analysed by looking at the difference between the known *K* and the number of estimated components and at the frequency of such number along the chain iterations. The number of estimated components was considered as the most popular number along the MCMC steps. Figure 1 shows that for *α*_*sim*_=1 the HDMM driven by *α*_*HDMM*_=1 seemingly converges to *K* uniformally along the average number of observations. This is only true for *α*_*HDMM*_=1*/M* when the average number is greater than four for *K* = 5 and greater than seven for *K* = 10. Under *α*_*sim*_=1*/M*, though, the number of estimated components seems to linearly increment along the average number of observations for both *α*_*HDMM*_ and is always higher when *α*_*HDMM*_=1*/M*. These results are complemented by Figure 2 that reports how frequent the estimated components are along the fit. When *α*_*sim*_=1 the frequency raises quickly for increasing observations in the case of *α*_*HDMM*_=1 but, only after a certain threshold in the other case *α*_*HDMM*_=1*/M*. Besides, with *K*=10 the frequency slowly increases for *α*_*HDMM*_=1*/M*. A different scenario is portrayed by the test on *α*_*sim*_=1*/M*, where frequency slowly drops for higher average number of observations but it is still higher than many cases w.r.t. *α*_*sim*_=1. Through all tests results modelled by *α*_*HDMM*_=1 show to achieve higher frequencies w.r.t. *α*_*HDMM*_=1*/M*, although only marginally if *α*_*sim*_=1*/M*.

**Fig 1.**
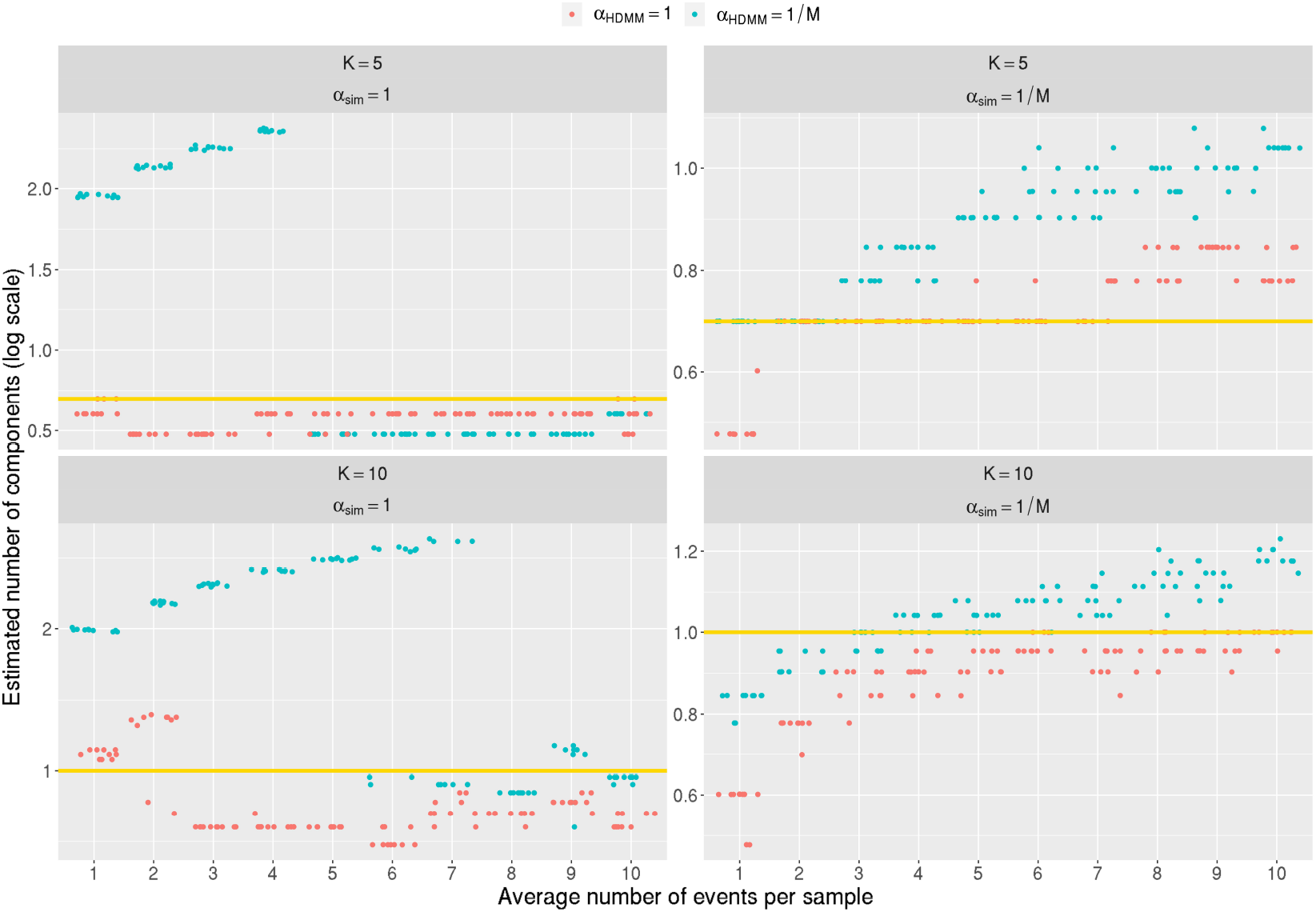
Number of components estimated by the HDMM (y-axis) is illustrated along the average number of observations per patient for several settings (x-axis). Logarithmic scale is followed on the y-axis and the yellow horizontal line stands at the expected log(*K*). Colors represent the different concentration parameters employed by the HDMM.

**Fig 2.**
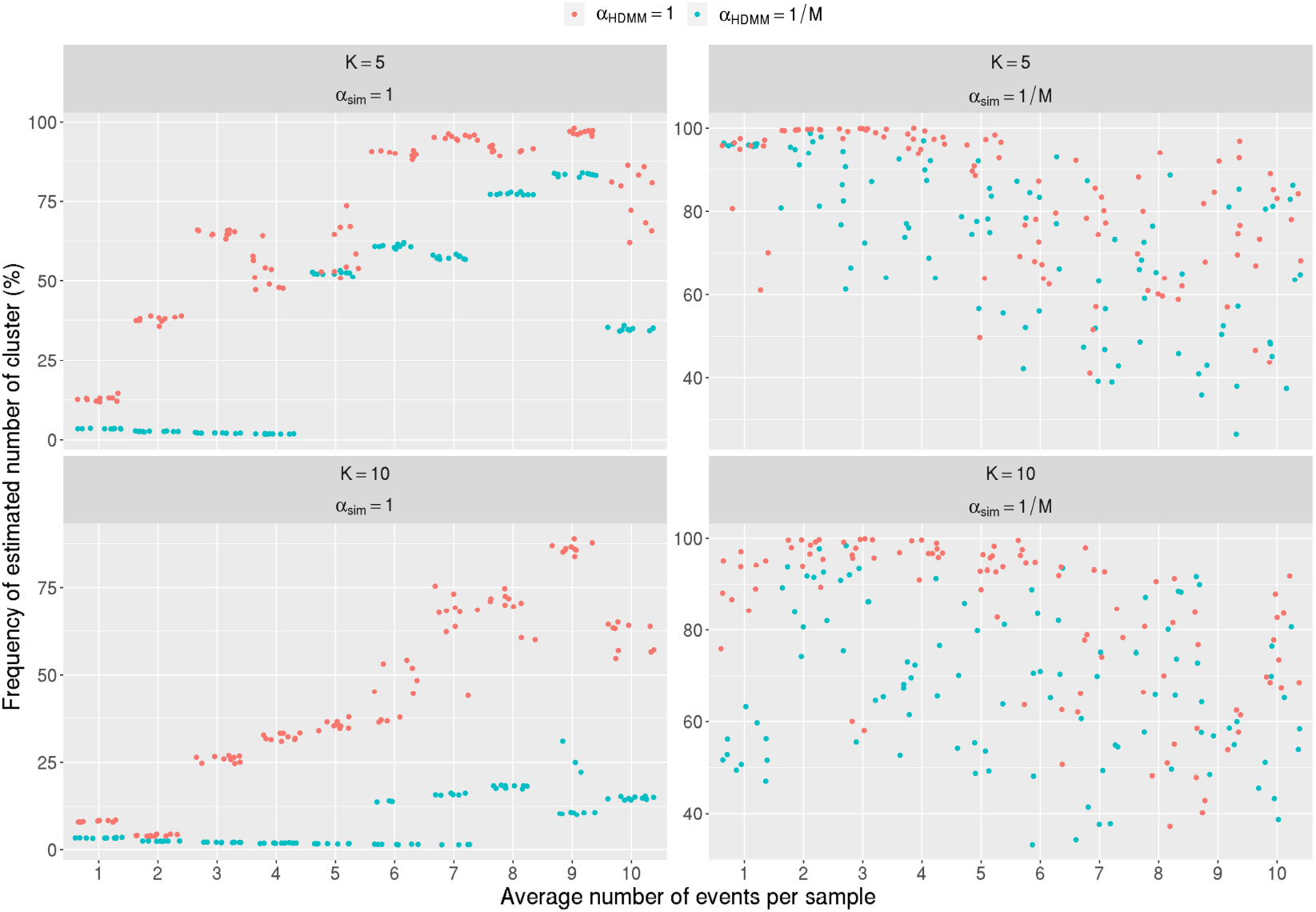
Dominance of the number of components estimated by the HDMM (y-axis) is reported over the average number of observations per patient (x-axis). The dominance is measured by the frequency of number occurence along the step of the MCMC used to fit the HDMM.Colors represent the different concentration parameters employed by the HDMM.

### Clustering patients with an automatic statistical approach

As introduced above, the outcome of a HDMM renders a matrix counting the number of times an observation is assigned to a component. If these counts are used to estimate for each component the parameters of a multinomial distribution, then the probability mass function (p.m.f.) of the multinomial can be used to cluster data samples entirely. Since the true component is known for each sample, then the result of the automatic clustering based on the multinomial p.m.f. can be quantitatively measured. Using the Adjusted Random Index (ARI) [20] as metrics for accuracy, the herein proposed approach achieves median performances roughly above 0.60 for all four combinations of *α*_*sim*_ and *α*_*HDMM*_ over all tested average number of observations for *K*=5 (Figure 3). This is not true for *K*=10 where performances are maintained high only for *α*_*sim*_ = 1*/M* (median ARI above 0.79) but drop around 0.3 for *α*_*sim*_ = 1. Besides, also for the *K*=5 the best ARI values resulted when *α*_*sim*_ = 1*/M*.

**Fig 3.**
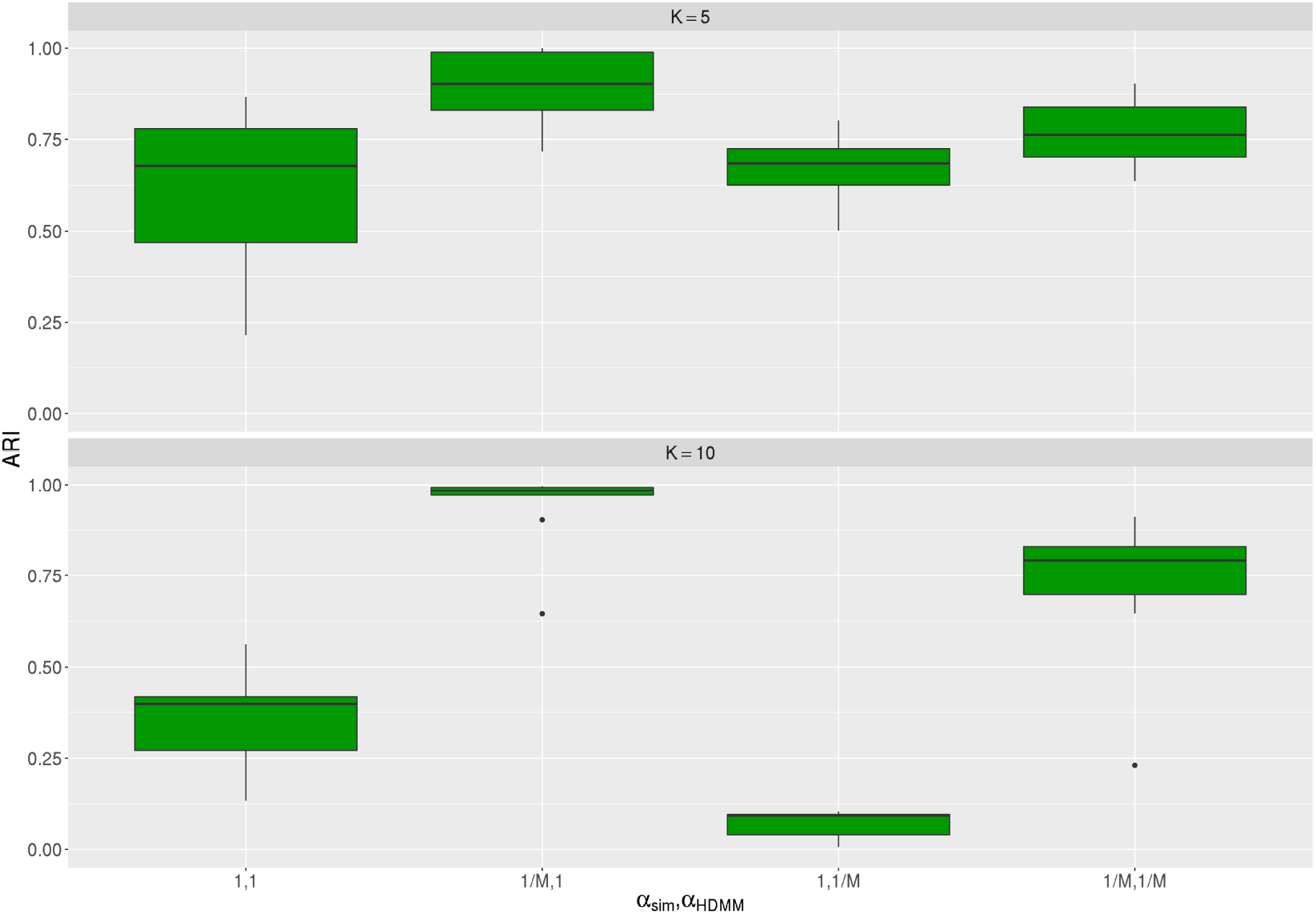
Adjusted Rand Index (ARI) of the novel multinomial-based clustering approach over all combinations of concentration parameters. This metric was adopted to compare two clusterings (i.e., the one initially simulated and the novel one) potentially using a different number of clusters.

### Advantages of using MFNCH distributions

Instead of considering the components as multinomial distributions they can be modelled as initially simulated, i.e., as MFNCH distributions. This alternative way also brings a new option to perform data sample clustering when it is feasible to compute the p.m.f. of the MFNCH distribution, which strongly depends on the ability to calculate the partition function of the distribution. In our test, the MFNCH p.m.f. was calculated for all detected components and the samples were clustered automatically. Roughly, the performance of the MFNCH-based clustering is never greatly lower than the alternative multinomial-based approach (Figure 4). In fact, the measured negative ARI gap of this clustering is always lower than 0.1. Though, additional uplift of ARI can occur and this is particularly typical when HDMM with *α*_*HDMM*_ = 1*/M* is employed to fit simulated data being generated using *α*_*sim*_ = 1*/M*. In such situation the increase in ARI was on average 0.1 and at most was greater than 0.2.

**Fig 4.**
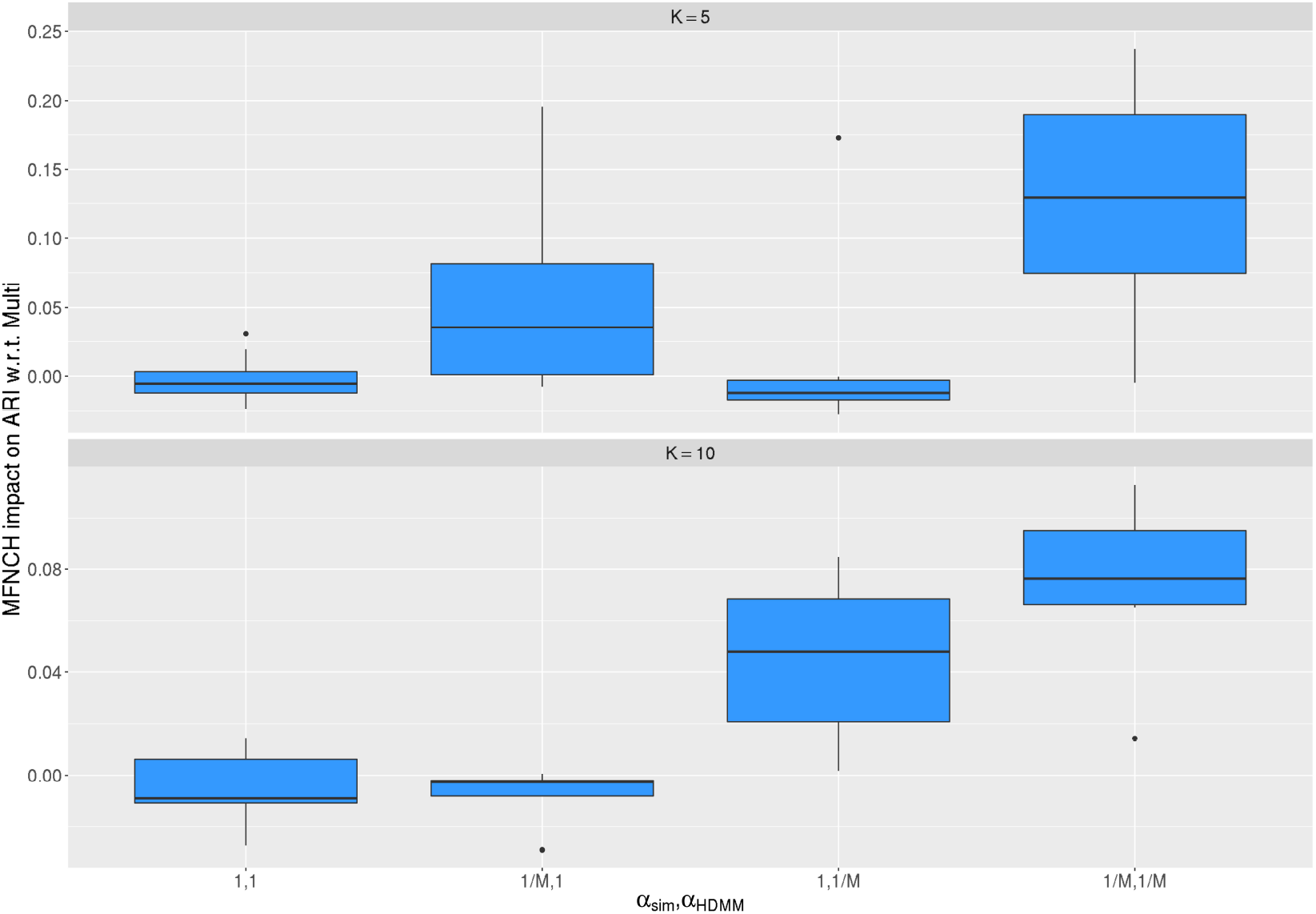
Variation in ARI when clustering is performed using p.m.f. of MFNCH distributions.

### Real data testing

To showcase the potential impact of using either the multinomial or MFNCH based approaches for clustering, a sample from a real fitted HDMM was taken into account. This HDMM was used to model the genomic data of more than one thousand Acute Myeloid Leukemia (AML) patients and was eventually utilized to propose a new clinical disease classification. Table 1 shows previously-reported genetically-defined AML types along with their most frequent genomic alterations. It should be noted that the frequency of the genomic alterations was calculated only after patients clustering. Table 2 compares which genomic alterations are prioritized when the components are estimated either as multinomials or as MNFCH distributions before clustering. In the multinomial scenario, the top five most probable genomic alterations are reported; whereas, in the MFNCH case, the top five most important alterations are reported. Importantly, the sixth component was manually separated in previous work for medical reasons and was maintained here to preserve methodological consistency with the HDMM perspective. The multinomial and MFNCH prioritize alterations differently. In MFNCH, mutations previously reported as very relevant such as *ASXL1* and *RUNX1* disappear. The *MLL* mutation changes the category and *FLT3*^*ITD*^ decrease in rank. The bold-marked alterations in the Tables were the discriminant used in previous work to perform patients clustering. The MFNCH-based approach shows to always bring on top such discriminant, whereas the multinomial-based approach achieved the same only in seven out of ten components.

**Table 1.** Genomic subgroups from real HDMM execution on AML dataset estimated as reported in literature. Bold genomic alterations indicate the main genomic discriminant used to perform patients clustering.

|  | Genomic component | Reported frequent alterations |
| --- | --- | --- |
| 1 | AML with *NPM1* mutation | ***NPM1***, *DNMT3A*, *FLT3<sup>ITD</sup>*, *NRAS*, *TET2*, *PTPN11* |
| 2 | AML with mutated chromatin, RNA-splicing genes, or both | ***RUNX1***, ***MLL***, ***SRSF2***, *DNMT3A*, ***ASXL1***, ***STAG2***, *NRAS*, *TET2*, *FLT3<sup>ITD</sup>* |
| 3 | AML with *TP53* mutations, chromosomal aneuploidy, or both | **Complex karyotype**, **-5/5q**, **-7/7q**, ***TP53***, **-17/17p**, **-12/12p**, +8/8q |
| 4 | AML with inv(16) | **inv(16)**, *NRAS*, +8/8q, +22, *KIT*, *FLT3<sup>ITD</sup>* |
| 5 | AML with biallelic *CEBPA* mutations | ***CEBPA<sup>biallelic</sup>***, *NRAS*, *WT1*, *GATA2* |
| 6 | AML with t(15;17) plus AML with t(6;9) | **t(15;17)**, *FLT3<sup>ITD</sup>*, *WT1* plus t(6;9), *FLT3<sup>ITD</sup>*, *KRAS* |
| 7 | AML with t(8;21) | **t(8;21)**, *KIT*, -Y, -9q |
| 8 | AML with *MLL* fusion genes | **t(x;11q23)**, *NRAS* |
| 9 | AML with inv(3) | **inv(3)**, -7, *KRAS*, *NRAS*, *PTPN11*, *ETV6*, *PHF6*, *SF3B1* |
| 10 | AML with *IDH2<sup>R172</sup>* | ***IDH2<sup>R172</sup>***, *DNMT3A*, +8/8q |

**Table 2.** Prioritization of genomic alterations in the genomic subgroups when HDMM components are considered both as multinomials and as MFNCH distributions. Bold genomic alterations indicate the main genomic discriminant used to perform patients clustering.

|  | Multinomial prioritization | MFNCH prioritization |
| --- | --- | --- |
| 1 | ***NPM1***, *DNMT3A*, *FLT3<sup>ITD</sup>*, *TET2*, *NRAS* | ***NPM1***, *MYC*, *MLL5*, *DNMT3A*, *RAD21* |
| 2 | ***RUNX1***, ***SFRS2***, ***STAG2***, ***MLL***, ***ASXL1*** | ***SFRS2***, ***STAG2***, *ZRSR2*, *CUX1*, *MPL* |
| 3 | **Complex karyotype**, **-5/5q**, ***TP53***, **-17/17p**, -7 | **Complex karyotype**, **-5/5q**, ***TP53***, **-17/17p**, abn(3q) |
| 4 | **inv(16)**, *NRAS*, *FLT3<sup>TKD</sup>*, +8/8q, +22, | **inv(16)**, +22, *KIT*, *NRAS*, -7q |
| 5 | ***CEBPA<sup>biallelic</sup>***, *GATA2*, *NRAS*, *WT1*, *CEBPA<sup>monoallelic</sup>* | ***CEBPA<sup>biallelic</sup>***, *GATA2*, *EP300*, *FBXW7*, *WT1* |
| 6 | *FLT3<sup>ITD</sup>*, **t(15;17)**, *WT1*, +8/8q, *FLT3<sup>TKD</sup>* | **t(15;17)**, **t(6;9)**, *WT1*, *FLT3<sup>ITD</sup>*, +8/8q |
| 7 | **t(8;21)**, -Y, *KIT*, -9q, *KDM6A* | **t(8;21)**, -Y, *KDM6A*, *PRPF40B*, *MLL3* |
| 8 | +8/8q, **t(x;11q23)**, -4/4q, Complex karyotype, +11/11q | **t(x;11q23)**, +8/8q, -4/4q, *MPL*, *MLL5* |
| 9 | -7, *NRAS*, **inv(3)**, *PTPN11*, *KRAS* | **inv(3)**, *ETV6*, *SF3B1*, -7, *KRAS* |
| 10 | ***IDH2<sup>R172</sup>***, *DNMT3A*, +8/8q, *CREBBP*, -7q | ***IDH2<sup>R172</sup>***, *CREBBP*, -7q, *BCOR*, *DNMT3A* |

## Discussion

In this work the performances of HDMMs on binary simulated data, which is close to the usual genomic data from onco-hematology, were examined in-depth to optimize their employment. To simulate binary data and to understand how they can be modelled using HDMMs some key considerations were made. Frequently, binary data are viewed as count data and, as such, they are modelled by multinomials. Although a vector of binary values can be sampled from multinomials, sampling from multinomials does not guarantee to yield a vector of binary values. In fact, the multinomial distribution models samplings with replacements, i.e., an observation can be drawn many times, and its parameters are probabilities, which exactly expresses that some observations are more probable than others. To address this issue, the simplest solution is to assume that a binary vector comes from a sampling without replacement. This case is modelled by the well-known multivariate hypergeometric distribution. Since the parameters of the multivariate hypergeometric distribution are the number of times an observation can be sampled, if all parameters are set to one, the domain of the distribution is only composed by binary data. That is, the multivariate hypergeometric distribution can be tailored to produce only binary data. Though, differently from multinomials, the multivariate hypergeometric distribution, with all its parameters set to one, does not provide ways to prioritize observations, which means that only one hypergeometric distribution exists. Since this would prevent the usage of a mixture model, an intuitive solution would be to exploit a distribution modelling sampling without replacement but with prioritized observations. This case falls under the domain of the so-called multivariate non-central hypergeometric (MNCH) distributions that endow each observation of the multivariate hypergeometric distribution with an additional parameter, i.e., a weight. The larger the weight of an observation, the higher it is the bias of its occurrence. Therefore, in this work, data were simulated from a MNCH distribution, in particular from the one formulated by Fisher (MFNCH). To randomly determine MFNCH distributions, their weights were sampled as the parameters of multinomials from a Dirichlet distribution. After deciding what distribution better fits the nature of the desired binary data, several mixtures of MFNCH distributions were generated to cover many possible scenarios based on three variables. The first variable was the number of distributions underlying the mixture and two cases were taken into account, i.e., *K* = 5, 10. The second variable was used to set all concentration parameters of the a priori Dirichlet distribution and two extreme situations were studied: *α*_*sim*_ = 1 and *α*_*sim*_ = 1*/M*, where M stands for the size of observations (i.e., number of features). The former yields MFNCH distributions with almost uniform weights, which entails that the distributions of the mixture tend to be similar. The latter, contrarily, rendered MFNCH distributions with a few high weighted and lots low weighted observations, which increased the difference heterogeneity in the mixture. Third, the average number of total observations per sample in the simulated data was ranged from one to ten. This short low range was chosen to be in the order of magnitude of the number of average mutations per patient of onco-hematological datasets, hardly greater than five with the usual gene panels. Following data simulation, a HDMM of multinomials was fitted. Since data were actually generated by MFNCH distributions, using these distributions in the mixture in place of multinomials was taken into account. The reasons why multinomials eventually remained the underlying distribution of the mixture were motivated both by the actual state-of-the-art in onco-hematological studies but also by the time-consuming task of computing the partition functions of the MFNCH distributions that makes their implementation slow and hardly scalable. Since HDMMs were also provided with a concentration parameter *α*_*HDMM*_, the influence of this variable was additionally observed as for *α*_*sim*_, i.e. either *α*_*HDMM*_ = 1 or *α*_*HDMM*_ = 1*/M*. In fact, when *α*_*HDMM*_ = 1 the HDMM favors the process to retrieve uniform-like distributions. In contrast, when *α*_*HDMM*_ = 1*/M* the HDMM searches for low-overlapping distributions. The combinations of all four variables allowed to dissert the modelling of the data from different standpoints.

Initially, the convergence of the HDMMs was assessed observing the MCMC behind the fitting procedure. Along the MCMCs, the number of estimated components changes as well as their estimated parameters but, upon ideal convergence, such number and parameters should stabilize. That is, the statistical fluctuations along the chains should incrementally decrease. Besides, the perfect result is accomplished when the number of components coincides with the expected K number of distributions. Therefore, the most frequent number of components over 10^4^ MCMC iterations was considered as the best estimate for K and its frequency quantified its statistical fluctuations. The results showed different convergence behaviors along the average number of observations per sample between expected distributions simulated with *α*_*sim*_ = 1 and with *α*_*sim*_ = 1*/M*. In the first case, when simulated distributions have similar and uniform weights, higher average number improves convergence, especially for *α*_*HDMM*_ = 1. Interestingly, if *α*_*HDMM*_ = 1*/M*, the number of components increases and its frequency is low until a certain average number of observations per sample, between five and seven, where both metrics abruptly improve. These results highlight that many similar expected distributions are better recognized after a certain average number of total observations. Differently, when *α*_*sim*_ = 1*/M*, both *α*_*HDMM*_ values indicate that lower the average of total observations results in a better convergence. Intuitively, this scenario suggests that when expected distributions are extremely concentrated on a few observations they are well recognizable even given a single observation. When, instead, these distributions are sampled many times, low weighted observations are drawn and add noise to the HDMM fitting process. These results on convergence express two main concepts: the scenario with *α*_*HDMM*_ = 1 is more likely to converge but it is more conservative, i.e., little number of components; the scenario with *α*_*HDMM*_ = 1*/M* does not always converge (even lower than 50%), but when it does, it may in-depth describe the expected distributions (with a risk of overfitting).

As explained above, the main outcome of the HDMM is a matrix counting the number of times an observation is assigned to a component. Although the MCMC performs clustering, it clusters observations and not entire samples. This renders a landscape of co-occurrence between observations but it leaves open the possibilities to assign samples. In onco-hematology, this outcome is usually implemented with clinical knowledge to define new classifications so that patients are clustered based on a few driver observations. Though, the statistical framework of the HDMM already provides a totally unbiased approach to assign patients. In fact, since the fitted HDMM consisted of a mixture of multinomials, such multinomials may be estimated from the outcome matrix. Namely, the observation counts for each component can be normalized to one to approximate the associated multinomial. It follows that a sample can then be assigned to the component with the largest probability of sampling it. Probabilities are intuitively calculated using the multinomial probability mass function (p.m.f.). By doing so, samples are clustered based on all their observations and not only on a few driver ones. The ARI between expected and observed components, which shows how strongly two clustering results correlate, was chosen to quantify accuracy because the number of expected and observed components may differ. Noteworthy, the best performances are achieved when distributions were simulated with *α*_*sim*_ = 1*/M*, which makes sense since the distributions differ the most. In contrast, when they are similar and uniform (*α*_*sim*_ = 1) ARI metrics drop and seem to decrease additionally when K is larger. This may still be explained by the lack of strong discriminants between the simulated HDMM distributions whose multinomials were conditioned by concentration parameters equal to one (*α*_*sim*_ = 1). In other words, strong different patterns can be captured by the HDMM and their samples are accurately assigned by an intuitive statistical approach based on the usage of the multinomial probability mass function. To our knowledge this approach was never utilized in onco-hematology.

To further stress how such statistical clustering approach can be further refined, the outcome matrix of the HDMM was also used to estimate each component as a MFNCH distribution. This subtle refinement step was motivated by the following rationale. If an observation is very popular across the components it tends to have a prominent role in all of them w.r.t. to the other observations. Now, in case of an unpopular observation, i.e., rarely emerging, is completely assigned by the HDMM to a single component that is also characterized by some popular observations. This univocal assignment remains unseen when a multinomial is calculated, since the normalization gives higher probabilities to popular observations rather than prioritizing the unpopular but extremely discriminant observations. To address this possible oversight in prioritizing observations in components, the components were modelled as MFNCH distributions whose weights were approximated given the outcome matrix of the HDMM and the observations sizes, which are equal to the number of times observations occur in all simulated data. In analogy with the multinomial case, the p.m.f. of the MFNCH distribution was employed to cluster samples. The results interestingly showed that the previous ARI metrics are approximately at least as accurate as the ones obtained by the multinomial-based approach. Besides, a clear boost is achieved in the scenario with *α*_*sim*_ = *α*_*HDMM*_ = 1*/M*. When a mixture of very well characterized MFNCH distributions generate data (*α*_*sim*_ = 1*/M*) and are modelled by a HDMM driven by a low concentration parameter, i.e., searching for highly observation concentrated components (*α*_*HDMM*_ = 1*/M*), using MFNCH to prioritize observations within components and cluster sample is significantly beneficial.

To appreciate the difference on real data, the final outcome of a HDMM of multinomials fitting a dataset of Acute Myeloid Leukemia (AML) patients was compared on a published genetical and clinical dataset. The difference between the prioritized genomic alterations becomes clear in almost all genomic components when they are estimated as multinomials or as MFNCH distributions. The enhanced correspondence with the clinically oriented components when MFNCH is employed suggests that patients may be also clustered using the p.m.f. of the MFNCHs. While in large parts consistent with published findings, this alternative clustering prioritizes distinct genetic abnormalities hence generating hypotheses for subsequent studies that focus on the role of distinct genes in the pathology of AML. In particular, the downranking of the *ASXL1* and *RUNX1* mutations appears of interest and highlights the potential of the new MFNCH approach in hematologic malignancies. It is worthy to note that this approach is easily scalable and can be generalized to new patients.

## Conclusion

This work contributes to shed light on the consistent and optimized usage of HDMM on binary data and extend its potentiality thanks to an intuitive and scalable way of clustering samples. Additionally, it shows how multinomials can be outperformed by MFNCH in terms of clustering and interpretation, as portrayed by the example on a public AML dataset. New classification systems in onco-hematology may be inspired and motivated by the novel statistical clustering approach reported herein, which may be further tailored by clinical experts to improve patients stratification and ultimately personalized treatments.

## Author contributions statement

DD conceptualized and designed the study, conducted analysis, drafted the manuscript; ES, JME, LC provided data curation; ATT, JMT, MGDP provided clinical supervision; MB, RSR provided study materials; TM, AM, CS performed data analyses; CS, EG supervised data analysis and interpretation; JMHR, LB provided administration support; GC designed and coordinated analyses, plus supported interpretation.

## Data availability statement

All data and code is available on a GitHub repository at https://github.com/DanieleDallOlio/HDMM_refinement/tree/main.

## Acknowledgments

The authors thanks the HARMONY Healthcare Alliance Consortium: Alberto Hernández Sánchez, Angela Villaverde Ramiro, Caroline A Heckman, Axel Benner, Jurjen Versluis, María Abáigar, Marta Anna Sobas, Peter JM Valk, Klaus Metzeler, Teresa González, Joaqúin Martínez-López, Marta Pratcorona, Frederik Damm, Ken I Mills, Christian Thiede, Maria Teresa Voso, Guillermo Sanz, Konstanze Döhner, Michael Heuser, Torsten Haferlach, Rubén Villoria Medina, Michel van Speybroeck, John E Butler, Brian James Patrick Huntly, Gert Ossenkoppele, Hartmut Döhner.

